# BioJS-HGV Viewer: Genetic Variation Visualizer

**DOI:** 10.1101/032573

**Authors:** Saket Choudhary, Leyla Garcia, Andrew Nightingale, Maria Martin

## Abstract

**Motivation:** Studying the pattern of genetic variants is a primary step in deciphering the basis of biological diversity, identifying key ‘driver variants’ that affect disease states and evolution of a species. Catalogs of genetic variants contain vast numbers of variants and are growing at an exponential rate, but lack an interactive exploratory interface.

**Results:** We present BioJS-HGV Viewer, a BioJS component to represent and visualize genetic variants pooled from different sources. The tool displays sequences and variants at different levels of detail, facilitating representation of variant sites and annotations in a user friendly and interactive manner.

**Availability:** Source code for BioJS-HGV Viewer is available at: https://github.com/saketkc/biojs-genetic-variation-viewer

A demo is available at: http://saketkc.github.io/biojs-genetic-variation-viewer

## 1 INTRODUCTION

With the advent of next-generation sequencing technologies, it has been possible to profile genomes in larger numbers. One of the chief outcomes of such projects has been the cataloging of genetic variants in database resources such as dbSNP (Smigielski *et al*. (2000)) and Catalogue Of Somatic Mutations In Cancer (COSMIC) (Forbes *et al*. (2011)). These catalogs contain sets of genetic variants found in species such as mouse, rat, zebrafish and human which can be utilized to study evolutionary relationships, population diversity and disease specific variations. The COSMIC database is a curated set of somatic mutations as observed in cancer samples. COSMIC’s release *v70* contains 1,564,699 unique variants taken from 1,029,547 samples covering 28,735 genes. dbSNP(Sherry *et al*. (2001)) build 142 contains two orders of magnitude more human variants than COSMIC with 112 million refSNPs. The availability of such a large amount of data makes the analysis and interpretation much more challenging.

Any exploratory attempt at analyzing the variation data would involve visualizing variants across the genome to determine specific sites, if any, where the mutations are more frequent or are absent completely. Thus, visualization is critical for interpreting the vast catalogs of variants. There are various visualization tools available: COSMIC Genome Browser (Forbes *et al*. (2011)), for example, allows visualization of variant sites alongside the annotations. The UCSC genome browser (Karolchik *et al*. (2003)) is another widely used visualization browser. These tools also have a server side component and offer limited flexibility when dealing with style based changes. In particular, there is limited support for visualizing variants by applying various filters, which we discuss later.

BioJS-HGV Viewer is a BioJS (Gómez *et al*. (2013)) component developed to visualize genetic variants in a comprehensive manner. Using it requires very limited knowledge of HTML and javascript. BioJS is an open source javascript library providing various components to visualize biological data. The visualizations are web based and hence are almost platform independent.

## 2 METHODS

The functionality provided by BioJS-HGV Viewer has two mode views: (a) Overview (b) Detailed or Zoomed View

The core library of this component is designed to handle both DNA and protein variants. However, the current implementation has only been tested with protein variants. These variant sites have been generated by using a webservice provided by the Universal Protein Resource (UniProt). The webservice is still on a development stage and thus is not yet publicly available. This service has an indexed database of protein variants as reported in the COSMIC and UniProtKB (Consortium *et al*. (2014)) database and is made available as a JSON file, with support for standard data formats, such as the Variant Call Format or VCF, (Danecek *et al*. (2011)) being currently implemented.

The demo at http://saketkc.github.io/biojs-genetic-variation-viewer loads the variants for protein *J3KP33*, by default. The component however allows loading other proteins by passing an additional argument to the web address(URL). For example: http://saketkc.github.io/biojs-genetic-variation-viewer?q=P00533.

By default, SIFT and Polyphen consequence probability are averaged and the type of mutations are then decided based on this average score. The component however allows user to choose from either or all of the consequence probability.

The user can also choose to hide a particular category of mutations. Both the overview and detailed mode have another ‘expanded view’ where these mutations can further be separately visualized as *Stop Gained, Missense* and *Splice Region*.

All the visualizations are rendered as scalable vector graphics(SVG) using the *d3js* javascript library.

### 2.1 Overview Mode

In the default mode (Fig. 1) the viewer displays variant information in a condensed format using a stacked bar char displaying the number and *type of mutations* at each site. Detailed annotations are displayed on hovering over the rectangle as a *tooltip*.

The *type of mutations* are classified as: (a) Benign (b) Damaging (c) Mixed

The ‘Mixed’ category represents an **intermediate** state between damaging and benign.

**Fig. 1.**
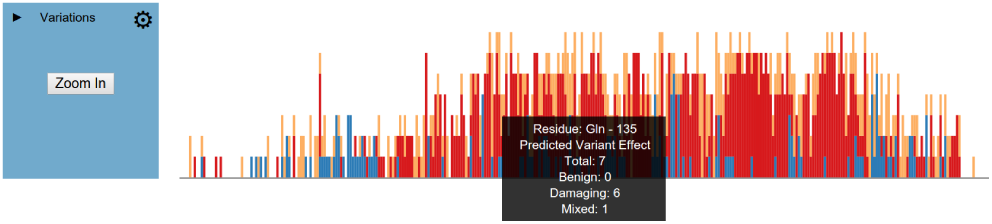
‘Overview’ of genetic variants as shown in by HGV viewer. Tooltips are used to display the number of mutations in benign, damaging and mixed categories.

The classification currently uses the predictions scores of Polyphen (Ramensky *et al*. (2002)) and SIFT (Kumar *et al*. (2009)). Polyphen scores are on a scale of [0, 1] with 1 indicating that the mutation is damaging and 0 indicating the mutation being benign. SIFT scores also operate on the scale of [0, 1] however 0 indicates a damaging mutation. The webservice has a database of all mutations across various proteins with pre-generated scores which can be retrieved as a JSON file.

The data thus received is parsed for calculating the number of mutations in each category. Each category is defined by threshold levels. For example a Polyphen score between 0.75 and 1.0 can be considered to reflect a damaging mutation. These threshold levels can be modified by the user. The height of each rectangular box depicting the mutation is dynamically adjusted based on the maximum number of variants at any site.

### 2.2 Detailed View

In the detailed view (Fig. 2) each individual amino acid on the protein is displayed as a rectangular box with all variants at that site, the height of the rectangle being proportional to the reported frequency. The box for variants is colored based on its type. On a *mouse over* action at the variant box, the tooltip shows detailed information about that particular mutation.

**Fig. 2.**
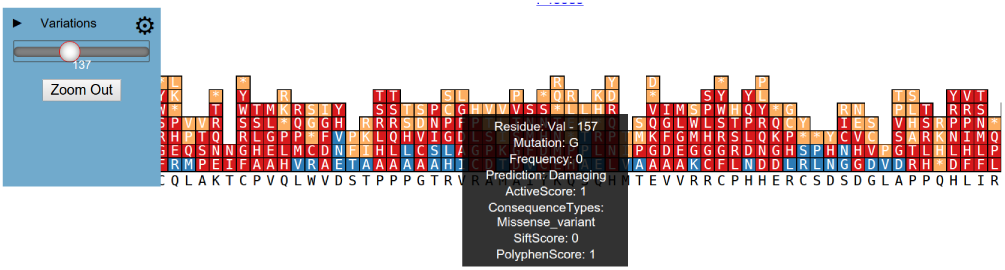
‘Detailed view’ of genetic variants. The SIFT/Polyphen scores and associated information with the mutations is rendered using tooltips

### 2.3 Expanded View

In both the ‘overview’ and ‘detailed’ view, it is possible to visualize the variants in separate individual views arranged as ‘Stop Gained’, ‘Missense’ and ‘Spliced Region’ variants. This provides a quick view of the distribution of different variant types. It is possible to easily switch between all the three views.

## 3 DISCUSSION

BioJS-HGV Viewer is an open source BioJS component that can be used as tool to visualize variants in a flexible manner. It can run entirely on the client side and hence can be used across various platforms.

The ability to visualize variants based on their categories *Benign, Damaging* or *Mixed* and apply further filters based on the mutation being *Stop Gained, Missense* or *Splice Region* is a unique feature.

The visualization provided by BioJS-Viewer can be the first step towards discovering more meaningful patterns in biological data. It can serve as a visual aid for studying the distribution of various types of mutations in DNA or amino acid sequences before performing detailed analysis.

## ACKNOWLEDGMENT

We thank the Google Summer of Code like and the BioJS community for supporting the original work.

*Funding:* S.C. was funded by the Google Summer of Code 2014 program sponsored by Google Inc.

